# Characterizing the speed and severity of mountain pine beetle spread under climate change with a mechanistic model

**DOI:** 10.1101/2024.06.27.601084

**Authors:** Micah Brush, Mark A. Lewis

## Abstract

In the last few decades, mountain pine beetle (MPB) have spread into Alberta, partially facilitated by climate change and warmer winters. Future effects of climate change on pine beetle spread are uncertain as warming is likely to affect both forest growth and beetle development. We here present a mechanistic model of pine beetle dynamics and characterize simulated outbreaks under climate change. This model includes key aspects of pine beetle biology, and we determine plausible ranges for each model parameter from available data. We then consider how forest growth, beetle brood size, and host resistance are likely to change in Alberta by the end of the century, and how this will relate to changes in model parameters. We simulate beetle outbreaks and quantify how the projected change in the distribution of the parameters will affect the period, speed, and severity of pine beetle spread. We find that simulated beetle outbreaks move more quickly than historically and are more severe.

## 1 Introduction

Forest pests are one of the most important drivers of forest disturbance, with impacts and management challenges expected to increase under climate change (Bale et al. 2002; Logan et al. 2003; Liebhold 2012). In particular, forest pest range expansion creates challenges for forest managers who are required to rapidly adapt to novel pest species (Ayres and Lombardero 2018). In addition to the ecological impacts, forest pests can lead to large economic costs from the loss or degradation of forests (Ayres and Lombardero 2000; Dhar et al. 2016; Corbett et al. 2016). Forest pests may also create feedback loops by increasing the number of forest fires or releasing carbon stored in forests (Ayres and Lombardero 2000; Kurz et al. 2008). Predicting the spread of forest pests under climate change is therefore important for managers to be able to more quickly respond to changing conditions and novel invasive species.

We here focus on mountain pine beetle (MPB, *Dendroctonus ponderosae* Hopkins), a destructive forest pest native to Western North America (Safranyik and Wilson 2006). In recent decades, MPB has undergone a climate facilitated range expansion to the North and East in Canada and into Alberta, with the expectation that MPB outbreaks may become more severe with climate change (Carroll et al. 2004; Cooke and Carroll 2017; Bentz et al. 2022; Janes et al. 2014; Sambaraju et al. 2012; Preisler et al. 2012; Cullingham et al. 2011; Cudmore et al. 2010; Logan and Powell 2001). Alberta in particular is expected to warm considerably, with every degree of global mean temperature increase corresponding to a two degree increase in mean winter temperature (Hayhoe and Stoner 2019; Eum et al. 2023). However, predicting how MPB spread will change in Canada over the next century is challenging given the different and varying effects of climate change on both hosts and beetles.

Climate warming is likely to have several effects on beetles themselves, given their development is temperature sensitive (Safranyik and Wilson 2006). In most parts of their range, MPB are univoltine, but in very cold areas they may take two years to develop (Bentz et al. 2014). Potentially the most important impact is that the northern range of MPB appears to be constrained by overwinter mortality of the larvae (Soderberg et al. 2021; Safranyik et al. 2010), which are able to survive to temperatures around -40C by generating cryoprotectants (Régnière and Bentz 2007; Lester and Irwin 2012; Rosenberger et al. 2017). In Alberta specifically, the number of days below -30C is expected to decrease substantially, especially in the north where a single degree of climate warming may result in over a week fewer days below -30C (Hayhoe and Stoner 2019). A reduction in the number of very cold days may therefore allow for further northward range expansion. In addition to a reduction of cold-induced mortality, beetle emergence is mediated by temperature (Bentz et al. 1991; Safranyik and Wilson 2006), and synchronous emergence is key to overcome host defenses (Raffa and Berryman 1983; Logan and Bentz 1999). Warmer temperature in some areas may actually decrease synchronous emergence (Logan and Powell 2004; Bentz et al. 2014; Bentz et al. 2022). In areas where they are currently semivoltine and require two years to mature, warmer temperatures may allow for single year development (Bentz et al. 2014; Bentz et al. 2022). There is additionally evidence that beetle dispersal is temperature sensitive (McCambridge 1971; Safranyik and Wilson 2006; Chen and Jackson 2017), and that beetles may be able to construct longer galleries at higher temperatures (Amman 1972).

Forest health is also likely to be impacted by climate change, though effects are uncertain. Lodgepole pine itself is expected to shift northward or to higher elevations, and by the end of the century may no longer have be well adapted for large regions of its current range, particularly in British Columbia (Hamann and Wang 2006; Coops and Waring 2011). In Alberta, hotter temperatures may also decrease the total area suitable for the growth of lodgepole pine (Monserud et al. 2008; Chhin et al. 2008a). However, in regions where lodgepole pine is still able to grow, it may be able to grow more quickly with longer growing seasons and warmer temperatures (Monserud et al. 2008). Seed selection and assisted migration may be able to mitigate the negative impacts on lodgepole pine growth to some extent and could increase pine productivity in the short term with only a few degrees of warming, but these strategies may become less effective under greater warming by the end of the century (Wang et al. 2006; Wang et al. 2010). In addition to temperature effects, precipitation patterns in Alberta are expected to change. The amount of precipitation may increase in the north of the province, but increases in temperature and the length of the growing season will likely lead to greater overall dryness (Hayhoe and Stoner 2019; Eum et al. 2023). The south of the province is unlikely to get more rainfall and so drought is likely to be more frequent (Bonsal et al. 2013; PaiMazumder et al. 2013; Hayhoe and Stoner 2019; Eum et al. 2023). Drought stressed trees are less able to produce the toxic resin that defends them against pine beetle attack (Safranyik and Wilson 2006), and so these changes in precipitation may increase overall forest susceptibility to MPB infestations even in areas that remain suitable for pine growth.

The complicated effects of climate change together with the long term nature of the changes makes models key to understanding impacts on forest health and MPB infestations. While statistical and machine learning models may do well on shorter time horizons, mechanistic models built from biological principals may be likely more able to capture the impacts of climate change as this requires prediction outside of historical norms. While there are existing models that study how pine beetle spread could be affected by climate change, they tend to be phenomenological (Cooke and Carroll 2017) or statistical (Sambaraju et al. 2012; Preisler et al. 2012) in nature.

We here apply a recently developed model (Brush and Lewis 2023; Brush and Lewis 2024) that links pine beetle dynamics and forest growth to study how pine beetle spread is likely to change in the next century in Alberta. This model is spatially explicit and mechanistic, with biologically interpretable parameters, and thus can make quantitative predictions about pine beetle spread. We identify four model parameters that are the most likely to be affected by climate change: the amount of time it takes the forest to grow, the survival fraction of juvenile trees, the host resistance, and the brood size of MPB. Changing these parameters allows us to consider the effects of climate change on both forest stands as well as MPB. We first set the range of parameter values according to historical spread, and then identify how they may change in the future. We then study how the frequency, severity, and speed of pine beetle outbreaks may be affected by the changes in these parameters under climate change.

## 2 Methods

### 2.1 Model summary

We here summarize the model described in Brush and Lewis (2024), which is a spatial extension of the model described in Brush and Lewis (2023). This mechanistic model was initially formulated by combining the pine beetle dynamics in Goodsman et al. (2016) and the forest growth model in Duncan et al. (2015). It includes forest growth as well as key aspects of pine beetle dynamics. In particular, the model has an Allee effect that arises from a threshold number of beetles needed to overcome host defenses. Beetles in the model aggregate to overcome these defenses, and we here assume this aggregation is near optimal at small scales, as in Brush and Lewis (2024). This is meant to describe the complex chemical signalling of MPB, which release both aggregation and deaggregation pheromones to regulate their attack densities (Raffa and Berryman 1983; Berryman et al. 1985; Safranyik and Wilson 2006). The forest itself grows according to an age structured model where juvenile trees are not attacked by pine beetle, new growth is light limited (for which there is evidence in stands with MPB (Amman 1977)), and trees have no mortality from causes other than pine beetle attacks.

Mathematically, this model is a set of age-structured integrodifference equations (Lutscher 2019). That is, the model is discrete in time and continuous in space. This model structure works well to describe mountain pine beetle dynamics given their annual life cycle and single summer dispersal and reproduction. We first describe the non-spatial model, which is derived and studied in Brush and Lewis (2023). Each variable in the model is indexed by the year *t*. We first define the forest structure. We divide the total number of juvenile trees *J*_*t*_ into *N* age classes, indexed by *i* as *j*_*i*,*t*_. Juvenile trees grow each year with survival fraction *s* and become susceptible after *N* years, where the number of susceptible trees in year *t* is *S*_*t*_. The number of female beetles emerging is *B*_*t*_, which is given by assuming that infested trees from the previous year *I*_*t*−1_ each produce *c* beetles. The mean number of emerging beetles per susceptible tree is *m*_*t*_ = *B*_*t*_*/S*_*t*_ = *cI*_*t*−1_*/S*_*t*_. Each year, the fraction of susceptible trees that avoid infestation by MPB is given by the function *F* (*m*_*t*_; *k, φ*) (alternatively, the fraction of successfully infested trees is given by 1 − *F* (*m*_*t*_; *k, φ*)). The function *F* was derived assuming beetles distribute themselves among susceptible trees according to a negative binomial with aggregation parameter *k*, and that a threshold number of beetles *φ* is needed to overcome host defenses. Mathematically, the fraction of susceptible trees that avoid infestation in a year is

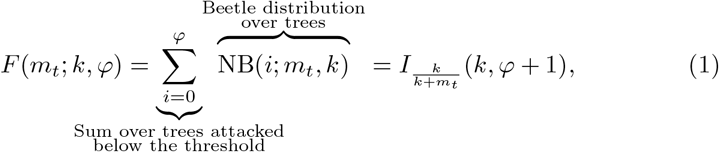

where 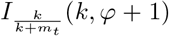 is the cumulative distribution of the negative binomial distribution, which is the regularized incomplete Beta function. We then assume that infested trees *I*_*t*_ block light to the forest floor for two years, where *I*_*t*−1_ gives the number of red top trees where the needles have turned red and *I*_*t*−2_ gives the number of gray snags where most needles have turned gray. To complete the forest growth model, we additionally assume that the total number of trees is fixed at *T* = *J*_*t*_ + *S*_*t*_ + *I*_*t*−1_ + *I*_*t*−2_. This means that new seedlings will grow wherever light is available, which occurs after juvenile mortality and when gray snags no longer block light to the forest floor.

We now follow the model extension to include space given by Brush and Lewis (2024). We index each variable by its spatial location *x*. We then assume the beetles disperse each summer according to a dispersal kernel *K*(*x* − *y*), which gives the probability that beetles at the spatial location *y* will disperse to the location *x*. We consider the model in one spatial dimension, which is equivalent to considering 2D radially symmetric dispersal with the appropriate choice of kernel. We note that while theoretically this model is continuous in space, in practice we will simulate this model with discretized space with cell size Δ*x*, where each spatial location will correspond to a few hundred square meters. The full evolution of the model is given by

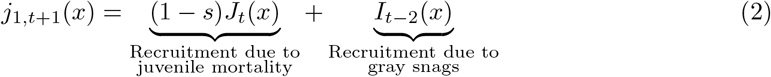

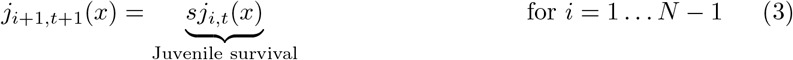

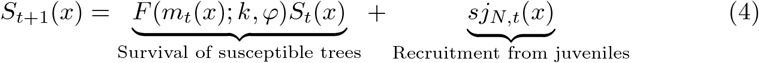

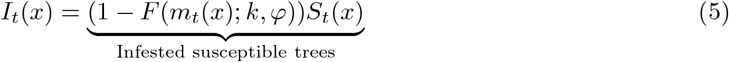

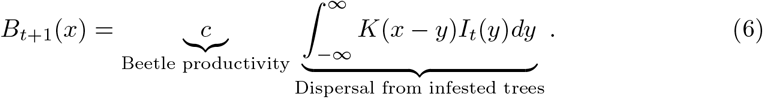

### 2.2 Beetle aggregation

As in the previous spatially explicit work with this model, we allow the aggregation parameter *k* to vary with the mean beetle density *m*_*t*_. This is because beetle aggregation is likely not constant over the course of an outbreak (Safranyik and Wilson 2006; Robertson et al. 2009; Boone et al. 2011; Chen and Walton 2011) or with stand density (Powell and Bentz 2014). We assume that beetles are excellent aggregators and so over the small spatial scale where they can detect pheromones, they aggregate optimally for each density. In practice, this means that we minimize the survival function *F* with respect to *k* for each value of *m*. We choose a minimum value of *k*_*min*_ = 0.01 for small *m*, and a maximum value of *k*_*max*_ = 100 for large *m*, as in Brush and Lewis (2024).

In addition to the minimum and maximum aggregation, we must choose a spatial scale for the aggregation. This will correspond numerically to the spatial scale of the grid, Δ*x*, as then within each grid cell beetles will aggregate optimally given the number of beetles within that cell. We will assume the beetles aggregate within a cell size of Δ*x* = 16 m (Brush and Lewis 2024), which is consistent with the assumptions in Powell and Bentz (2014), the aerial maps of Mitchell and Preisler (1991), and the results of Thistle et al. (2004) that find that the aggregation pheromones disperse to about 10% of their initial concentration within 10m. Note that our results are not very sensitive to this choice as long as is it a realistic value of tens of meters (Brush and Lewis 2024).

### 2.3 Beetle dispersal

Beetle dispersal is very complicated to model completely. Beetles can be lifted above the tree canopy and travel large distances (tens of kilometers) with the wind (Furniss and Furniss 1972; Jackson et al. 2008), but in general stay within a few hundred meters (Safranyik et al. 1992; Robertson et al. 2007). We again follow Brush and Lewis (2024) and assume a simple Laplace kernel for dispersal,

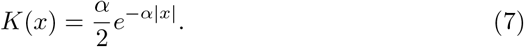

We note that the Laplace kernel in one dimension can be derived from a random walk dispersal with constant settling time model (Broadbent and Kendall 1953; Van Kirk 1995). While we will conduct simulations in 1D, we parameterize our kernel with 2D data and relate to the corresponding marginalized 1D kernel (Lewis et al. 2016).

In our simulations, we fix *α* = 0.001 m^−1^. We choose this value as in Brush and Lewis (2024), which predominately comes from analysis in Carroll et al. (2017), who use detailed survey data from Alberta to look at infestation spread at a larger scale. They find that about 75% of all new infestations occur within 2 km of the nearest previous infestation when averaged across all years from 2007 to 2013. This gives *α* = 0.0008 m^−1^, which we round up.

While there is some evidence that beetles may fly farther with warmer summer temperatures (Chen and Jackson 2017), assuming dispersal is fixed allows us to focus on the direct interaction between forest growth and resistance and the success of beetle infestations. Additionally, we note that a change in *α* simply corresponds to a change of units in our model, and so the speed scales roughly as 1*/α* (provided 1*/α* ≫ Δ*x*) (Brush and Lewis 2024).

### 2.4 Varying parameters under climate change

We now turn to the remaining parameters in our model that we will vary under climate change. These are *N*, the number of years it takes for seedlings to grow large enough to become susceptible, *s*, the fraction of juvenile trees surviving each year, *c*, the brood size or beetle productivity, and *φ*, the threshold number of beetles needed to overcome tree defenses. We obtain a reasonable range for each of these parameters from the literature, as in Brush and Lewis (2024). In that study, each parameter was fixed for simulations at a reasonable central value somewhere in the range. Here, we instead consider each parameter as a triangular distribution centered at its central value and with support over its range. Table 1 summarizes the range of present values and the references for how these ranges were obtained. See Brush and Lewis (2024) for additional details on how the present values of these parameters were obtained from the data.

**Table 1:** Parameter values and ranges estimated from the literature both in the historical range and under climate change.

| Parameter | Historical values |  | Projected values |  | Units |
| --- | --- | --- | --- | --- | --- |
|  | Central value | Range | Central value | Range |  |
| $N$, Years to become susceptible | 60 | 50-80 | 55 | 40-70 | Years |
| $s$, Juvenile survival | 0.99 | 0.98-0.998 | 0.994 | 0.99-0.998 | Survival/year |
| $c$, Brood size | 1300 | 400-6000 | 1700 | 1000-6000 | Beetles/tree |
| $\varphi$, Tree threshold | 250 | 150-600 | 250 | 150-320 | Beetles/tree |

We must additionally specify how each of these parameters are expected to change under climate change. We do not expect that changes in these parameters will be homogeneous in space. We consider how these parameters may change in Alberta, as this is the current leading edge of the pine beetle range and we wish to consider the risk of further spread across the boreal forest. In general, we expect lodgepole pine to be in its ideal climatic zone in fewer places in Alberta by the end of the century, but we do expect that there will still be areas in the north suitable for lodgepole pine (Monserud et al. 2008; Chhin et al. 2008a). In our simulations, we will assume that we are interested in a region still suitable for lodgepole pine growth. This allows us to study how MPB infestations are likely to change in regions where lodgepole pine is still being managed for forestry. We now consider each of these parameters individually and identify how they may change under climate change.

#### Years to become susceptible, *N*

The current range for this parameter is based on information and data from a large number of sources, with two methods. The first of which is taken directly from attack data (Safranyik 1968; Safranyik and Wilson 2006; Shore and Safranyik 1992). The second of which uses site index information combined with growth models to predict when trees reach a susceptible threshold. We use that same method here to predict this parameter under climate change, as there are available projections for lodgepole pine site index at the end of the century (Monserud et al. 2008). Under climate change, the area where lodgepole pine is well adapted could shrink dramatically by the end of the century (Monserud et al. 2008; Chhin et al. 2008a). However, in areas that are still suitable for lodgepole pine, the number of years it takes for trees to become susceptible is likely to decrease as a result of a longer growing season and warmer temperatures (Monserud et al. 2008). In some of these regions in Alberta, the site index (the average height a tree can reach after 50 years of growth above breast height) is projected to increase from 12-15 m to 21-24 m, with other regions at 18-21 m (Monserud et al. 2008). Using a height to dbh model (Huang 1999), we can obtain the approximate age where trees are about 25 cm in diameter, which we use as a threshold for susceptibility (Klein et al. 1978; Mitchell and Preisler 1991; Safranyik and Wilson 2006; Safranyik et al. 1974; Cole and Amman 1969). We then add the estimated 6 years for pine to reach breast height in Alberta (Huang et al. 1994; Thrower et al. 1994; Nigh and Love 1999). Finally, we assume a rate of 2 mm of radial growth per year (Chhin et al. 2008b; McLane et al. 2011; Heath and Alfaro 1990). With a site index of 21 m, the dbh at age 56 (the site index plus the number of years to breast height) can be calculated as 27 cm in the upper foothills and 25 cm in the lower foothills (Huang 1999), roughly our cutoff for susceptibility. We take this as our central value and consider site indices of 18 m and 24 m to obtain a plausible range for *N* . We use the lower foothills model with the assumption that under climate change, the upper foothills in the future will appear more like the lower foothills of the present. With this model, a site index of 18 m corresponds to a dbh of 19 cm and a site index of 24 m corresponds to a dbh of 32 cm. Assuming 2 mm of radial growth per year, we can then calculate the age to susceptibility as *N* = 56 + (25 − 19)/0.4 = 71 years for the site index of 18 m and *N* = 56 + (25 − 32)/0.4 = 38.5 years for the site index of 24 m. Thus, in our simulations we decrease the range of *N* to between 40 and 70 years with a central value at 55 years to represent that lodgepole pine grow more quickly in suitable areas.

#### Juvenile survival, *s*

The current range for this parameter is based on data from Brown and Navratil (1995), Bedford and Sutton (2000), and Rweyongeza et al. (2007). As with the number of years it takes to become susceptible, this is likely to vary strongly with region. In areas that are still suitable for lodgepole pine, this is unlikely to change dramatically but may increase slightly. However, in some regions, pine suitability will be much worse and we expect this to decrease as juvenile mortality will be much higher. In our simulations, we assume we are in an area that remains suitable for lodgepole pine, and so we increase the lower range of this parameter from 0.98 to 0.99 to align with data from a study site in BC where lodgepole pine is currently well adapted (Bedford and Sutton 2000). We additionally increase the central values to 0.994 to be intermediate between the lower and upper bounds.

#### Brood size, *c*

The current range for this parameter is based on data from Klein et al. (1978), Raffa and Berryman (1983), Goodsman et al. (2016), Safranyik (1968), and Cole and Amman (1969) and Safranyik and Wilson (2006). Note that in the case of both this parameter and the host threshold, we frequently scale data up from attacks per m^2^ to attacks per tree, using data from Klein et al. (1978) and estimates from Powell and Bentz (2009) and Strohm et al. (2013). In general, brood size is highly variable and depends on many factors other than climate, in particular the size of the host tree (Safranyik and Wilson 2006). However, one projected effect of warming is to decrease the overwinter mortality of pine beetle (Régnière and Bentz 2007; Cooke and Carroll 2017), particularly in northern Alberta where conditions remain suitable for lodgepole pine. Currently, cold winters with temperatures below about -40C kill overwintering larvae (Régnière and Bentz 2007; Lester and Irwin 2012; Rosenberger et al. 2017), and it is likely that the northern range of MPB is constrained by cold winter temperatures (Safranyik et al. 2010; Soderberg et al. 2021). Therefore, brood size is likely to increase as overwinter mortality decreases with warmer winter temperatures. In particular, the decrease in the number of very cold days that is expected in Alberta (Hayhoe and Stoner 2019; Eum et al. 2023) is likely to result in fewer winters with large mortality from cold. Here, we estimate this increase using the cold tolerance model included in BioSIM (Régnière and Bentz 2007; Régnière et al. 2014) for an area in northern Alberta near Willmore Wilderness Park (see Xie 2024). Under climate change (RCP 4.5), the average overwinter survival is projected to increase from 28% to 36%. We use this to adjust the central value of brood size by approximately 36*/*28 = 1.3 from 1300 beetles per tree to 1700 beetles per tree. In addition to the mean survival, the overall number of very cold winters where overwinter survival may be very low is important. We use data from Eum et al. (2023) on the cold spell duration index to adjust the lower end of the range of brood size. This metric is a measure of the consecutive days below the 10th percentile of the historical temperature. They find the current value of the cold spell duration index in Alberta is 3.2 days per year (Table S-3, Eum et al. 2023). This is projected to change to about 0.4 over all of Alberta by the end of the century under 3 degrees of warming (which corresponds approximately with RCP 4.5). This means a cold spell is approximately 8 times less likely. We decrease the odds of being below the historical central value of brood size of 1300 beetles per tree by 8 from 9/56 to 9/385, where these probabilities are calculated from the triangular distribution. This roughly corresponds to an increase in the lower bound from 400 beetles per tree to 1000 beetles per tree. Given that years with large outbreaks were not likely impacted by significant overwinter mortality in any case, we do not change the upper bound of the range from 6000 beetles per tree. Note that our increase in brood size here is similar to Cooke and Carroll (2017), who in their warming scenario increase the peak recruitment by a factor of two, but here we change the distribution rather than just the peak.

#### Tree threshold, *φ*

The current range for this parameter is based on data from Raffa and Berryman (1983), Waring and Pitman (1985), and Lewis et al. (2010) and Peterman (1974). In many of the areas suitable for lodgepole pine, this is likely to decrease under climate change due to drought stress that reduces the trees ability to defend themselves against pine beetle infestation (Safranyik and Wilson 2006; Cooke and Carroll 2017). While there is expected to be a moderate increase in precipitation, this is likely to be offset by longer growing seasons and hotter maximum temperatures (Hayhoe and Stoner 2019; Monserud et al. 2008). These effects are difficult to quantify as more frequent drought is likely to increase the frequency of years where trees are under drought stress, but this effect is most relevant in terms of the variance rather than the mean. Using the expected return period of extreme drought measured using the SPEI-12 metric (Table 8, Eum et al. 2023) we find that a small amount of warming (globally mean temperature increase of 1.5 degrees), the return period of extreme drought is approximately halved in many areas of Alberta. With 3 degrees of warming (which corresponds roughly to RCP 4.5) the return period of extreme drought decreases from 33 to 7 years in the Foothills area, which is the area of Alberta likely to be the most suitable for lodgepole pine. This corresponds to about 5 times more likely, and therefore we increase the risk of being below the central threshold of 250 beetles per tree by about 5 times. In this case, we hold the lower and central values fixed and decrease the upper threshold of resistance. The lower value of the threshold range likely comes from when trees are already drought stressed, and so we keep that number fixed at 150 beetles per tree. Additionally, the threshold of 250 is better estimated from data (Raffa and Berryman 1983), and so we keep that fixed. We decrease the upper threshold until the odds of being below the central threshold of 250 beetles per tree are increased by 5 times. This corresponds to a new upper bound of 320 beetles per tree, representing a decrease of in the number of years where trees are very able to defend against attack. Another source of evidence for this decrease is to consider the soil moisture index (SMI) from BioSIM data in the Willmore Wilderness region of Alberta (Régnière et al. 2014; Xie 2024). In this area of the foothills, the projected mean SMI under RCP 4.5 decreases from about 88 to 42, or by about a factor of 2, which is roughly in line with this change in the upper bound.

We have here identified plausible changes to each of the model parameters. While we cannot determine quantitatively exactly how these parameters will change, we expect the general magnitude and direction of these effects to be consistent with climate change in Alberta, and in particular in regions that are still suitable for lodgepole pine growth at the end of the century.

### 2.5 Simulation and outbreak metrics

We implement 1D spatial simulations as in Brush and Lewis (2024). We use a discrete grid with spacing Δ*x* = 16 m and use Fast Fourier Transforms to perform the convolution in Eq. 6. We interpret each discrete cell as an individual forest stand where beetles are able to aggregate optimally. We set each discrete cell to have total tree density *T* = 1, which allows us to interpret the values of *S*_*t*_(*x*), *j*_*i*,*t*_(*x*), and *I*_*t*−1_(*x*) or *I*_*t*−2_(*x*) as fractions of the total tree density in the stand at location *x*. We note that typical stands of lodgepole pine in Alberta have densities of a few thousand stems per ha (Baah-Acheamfour et al. 2023; Safranyik and Wilson 2006), and so with Δ*x* =16 m each cell contains approximately 50 to 500 trees.

As found previously, outbreaks from initial infestations in this model are characterized by large transient peaks as the infestation progresses (Brush and Lewis 2023). Over a very long time, these peaks settle to the equilibrium solution. It is also possible, depending on the parameter values, that initial infestations cannot spread or that the beetles die out (Brush and Lewis 2024). We will characterize simulated outbreaks using three metrics: the period between peaks, the size of the peaks, and the speed of the spread.

We conduct the numerical simulations as follows. We first randomly generate a set of parameters by drawing from triangular distributions for each of the parameters, either from their present distribution or their expected distribution under climate change. Figure 1 shows the present and projected triangular distribution of parameters generated from Table 1 from which we draw at random. We then check to see if an outbreak is possible given the random set of parameters. Mathematically, this means we check if the non-trivial equilibrium where 1 − *F* (*m*; *φ*) = *m/c* exists (Brush and Lewis 2023). If it does not, we set the outbreak period, size, and speed to 0 to indicate that the beetles do not outbreak. If it does, we set all variables to the left of the origin (*x* < 0) to their equilibrium values obtained by solving 1 − *F* (*m*; *φ*) = *m/c*. We set all trees to be susceptible to the right of the origin, or mathematically, we set *S*_0_(*x*) = 1 and *J*_0_(*x*) = *B*_0_(*x*) = *I*_0_(*x*) = 0 for all *x* ≥ 0. We assume that no trees were previously infested. With the initial conditions set, we then iterate the wave forward in time for some fixed number of steps.

**Fig. 1.**
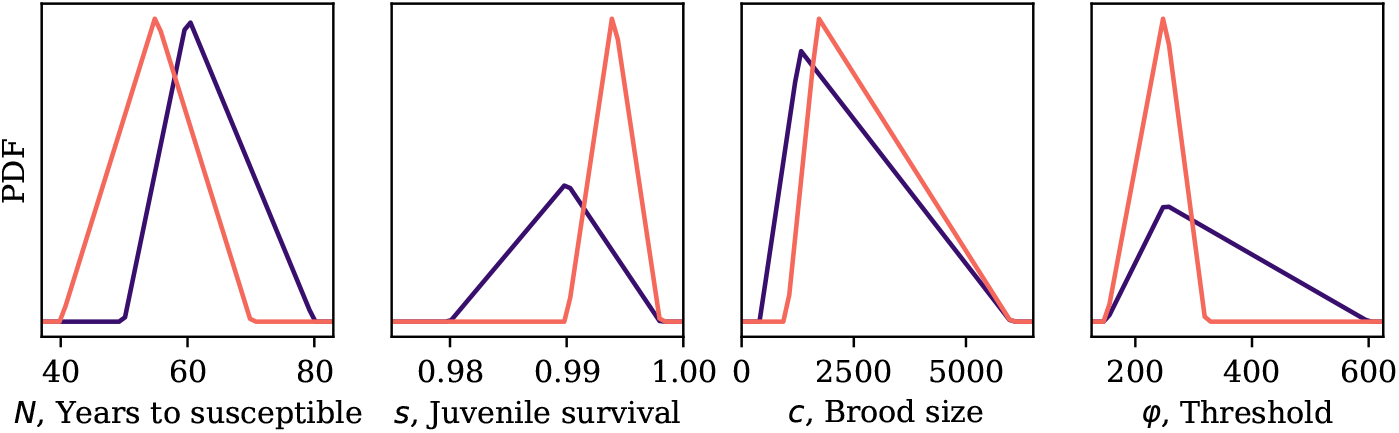
The triangular distributions from which we draw the four parameters that are altered under climate change. The parameters needed for these distributions are shown in Table 1. The present ranges are estimated from data as in Brush and Lewis (2024), and we then identify how we expect these distributions to be altered under climate change

Note, we choose this initial condition rather than setting the forest to the equilibrium density to the right of the equilibrium because this gives the asymptotic wave speed and size much sooner, which we will use to characterize the outbreak. Asymptotically, as the beetles spread to the right, the forest stands at the leading edge of the infestation will be composed entirely of susceptible trees. This is because the model does not include natural mortality and so stands will remain susceptible until they encounter beetles. Thus, setting the forest stands to the right of the origin to be composed only of susceptible trees initially means that we do not need as large of a grid to find the asymptotic wave speed.

To characterize the outbreak with a given set of parameters, we look for the first transient peak at *x* = 0. If that peak never occurs or is smaller than the equilibrium value of infestation, then we set the outbreak period, size, and speed to 0. If there is an outbreak, then at *x* = Δ*x*, one grid cell to the right of the origin, we do two numerical checks. First, we ensure we are able to calculate the size of the outbreak by checking that we have run the simulation for a long enough time that one and a half periods of iterations have passed since the initial transient peak at *x* = 0. Second, we check if the backwards wave that is generated at the boundary collides with the forward wave, and if it does, we consider the simulation only until the point where these waves collide. This ensures the boundary effects are not important for fast moving waves. We then calculate the wave metrics as follows.

#### Period of the wave

We calculate the time between the transient peaks by finding the peaks of the wave at the spatial location equal to the speed of the wave multiplied by *N* + 3, which was our previous estimate for the period of the outbreaks (Brush and Lewis 2024). This point was chosen to be away from the origin by roughly the distance the wave travels in one period to approximate the asymptotic period.

#### Size of the wave

We calculate the peak size of the infestation by averaging the peak number of infested trees over one entire period. We set the initial point in time by adding a half period to the time where the outbreak first occurs at *x* = Δ*x*.

#### Speed of the wave

We calculate the speed of the wave by finding the furthest point along the wave where the beetle population is greater than half of the equilibrium density at each time, which we call 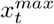. We then calculate 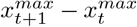 to determine the number of meters the infestation has spread in a year. In order to estimate the asymptotic wave speed, we average the last half of the simulation (or half the time until the wave collides with the backwards travelling wave from the boundary conditions).

## 3 Results

In order to characterize how these metrics are likely to change under climate change, we conduct 200 simulations with parameter values drawn from the distributions given by Table 1 and shown in Fig. 1, both with the present and projected distributions. We run each simulation for 400 time steps, and then calculate the period, size, and speed of the outbreak.

Figure 2 presents histograms for each of the three outbreak metrics. We find that the percent of simulations where an outbreak occurs increases from 85.5% to 99.5% of simulations under climate change. In general, the outbreaks shift to spreading more quickly and are more severe.

**Fig. 2.**
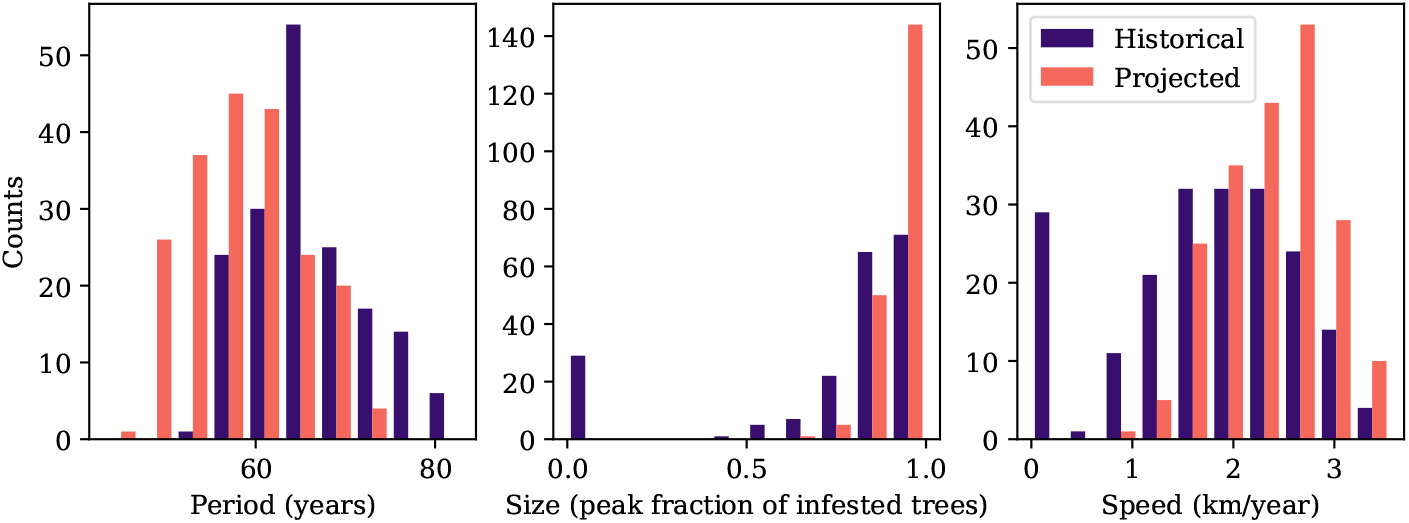
Histograms for the defined outbreak metrics under the present distributions of parameters and the projected distributions under climate change. The metrics describe the period, size, and speed of outbreaks. If the outbreak does not occur, the period, size, and speed are set to 0

The period between outbreaks also decreases as the pine forest grows more quickly. In all cases where an outbreak occurs, the period is equal to *N* + 3, as found previously in this model (Brush and Lewis 2023; Brush and Lewis 2024).

Therefore, the period of the outbreaks is completely determined by the time for the forest to regenerate in this model. On the other hand, the juvenile survival *s* does not appear to have a large effect on the metrics considered here within its biologically plausible range.

Previous analysis of this model (Brush and Lewis 2023; Brush and Lewis 2024) has demonstrated that the ratio *φ/c* is the parameter combination that largely determines the model dynamics. We call this combination the resistance per tree brood, and it acts as a kind of rescaling of the host resistance with respect to the number of beetles. Given this, we plot the size and speed of all simulations, including those that do not outbreak, against the ratio *φ/c* in Fig. 3. Here we can see that this ratio determines the speed of the outbreak and also appears to largely determines the peak size of the outbreak. This figure also shows more clearly the increase in the number of simulations where an outbreak occurs, where as we move to a smaller distribution of *φ/c* under climate change, the number of simulations where an outbreak can occur becomes larger. We note that once the beetles are able to spread, the size of the infestation increases sharply.

**Fig. 3.**
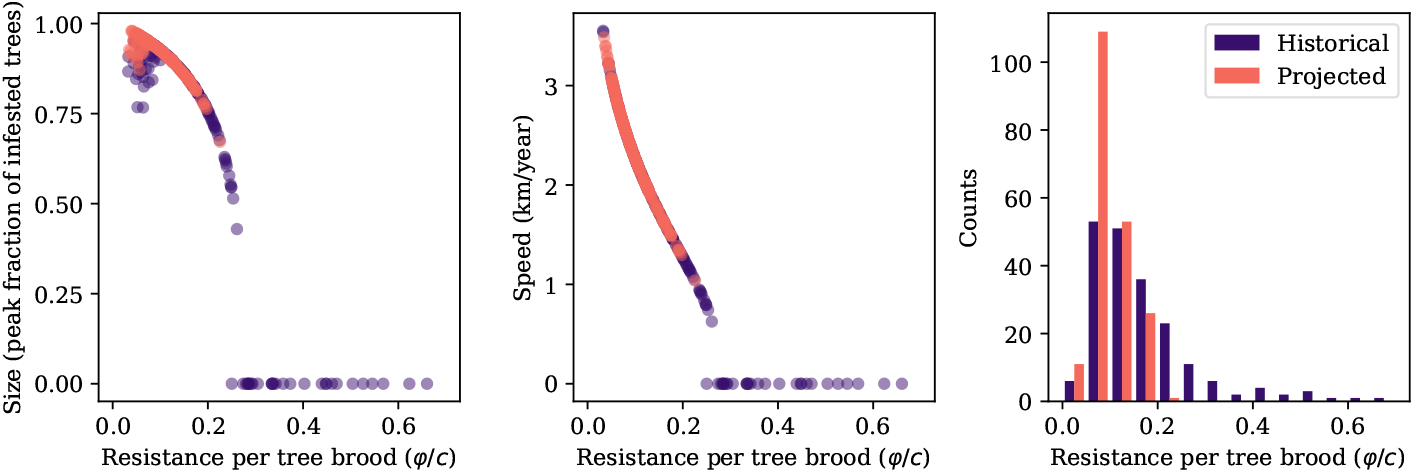
The size and speed of outbreaks under the present parameter ranges and those projected under climate change plotted against the parameter combination *φ/c*, and the distribution of *φ/c* under historical and projected conditions

## 4 Discussion

We have here investigated how the period, size, and speed of spread of mountain pine beetle may be altered under climate change using a carefully parameterized mechanistic model. We considered how the range each of the four parameters in the model are likely to change in regions still suitable for lodgepole pine in Alberta. We find that outbreaks are likely to occur in more stands, spread faster, and increase in severity.

We find that each of the four parameters will affect spread differently, with some more important than others. The return period of infestations is determined entirely by the number of years to become susceptible, *N*, and as has been found previously is equal to exactly *N* + 3 (Brush and Lewis 2023; Brush and Lewis 2024). The other parameter related to the forest growth model, the juvenile survival *s*, is unlikely to significantly affect any of the metrics of spread within the predicted range of the parameter. It is likely there is a small effect on the height of the second (and subsequent) transient outbreak peak, rather than on the peak fraction of infested trees overall, the metric considered here. This is because a small increase in juvenile survival can make a comparatively large difference for the number of juveniles that become susceptible over the time between outbreaks. With the central values historical values of *N* = 60 and *s* = 0.99, the second outbreak peak will only have about a fraction of approximately 0.99^60^ = 0.55 susceptible trees. With the central projected values of *N* = 55 and *s* = 0.994, this fraction increases to 0.994^55^ = 0.72, and so there are more susceptible trees for the beetles to consume on subsequent outbreaks with higher juvenile survival. However, note that these subsequent outbreaks and the juvenile survival after an outbreak depend strongly on forestry practices (Baah-Acheamfour et al. 2023). Additionally, in regions that are no longer suitable for lodgepole pine, this parameter may decrease more dramatically, leading to smaller, lower vigor pine trees that may not be as suitable of hosts for MPB. The ratio of the parameters related to the beetle dynamics themselves, the brood size *c* and the host tree threshold *φ*, largely determines the size and speed of spread (Fig. 3). This aligns with previous analyses with this model (Brush and Lewis 2023; Brush and Lewis 2024), but is interesting given how these parameters are likely to change under climate change. Given that the host threshold is likely to decrease on average under more severe drought conditions, and the brood size is likely to increase with warmer winter temperatures, this creates the possibility that stands previously resistant to MPB outbreaks may be able to support an outbreak. We can see this in Fig. 3, where the projected distribution of *φ/c* is below the threshold for an outbreak to occur. This also creates the possibility of more severe epidemics in places already susceptible, where the combination of drought and warmer weather may lead to beetles consuming more host trees.

We find that the size of the peak infestation is very high and the beetles consume nearly all the trees in many realizations of the parameters (Fig. 2). In reality, beetles may consume nearly all of the large diameter trees, but leave some trees above our supposed threshold of 25 cm dbh (Safranyik and Wilson 2006; Klein et al. 1978). We believe the explanation for this discrepancy is twofold, which highlight areas where our model is not perfectly capturing spread. First, the data that we are comparing to here are not exactly related to the distribution we are predicting, in that in our model the distribution is the complete variation of the parameters of *φ* and *c*. The variation over the full range of these parameters is unlikely to be realized in any single outbreak event, and it is more likely that the range of *φ/c* over a single outbreak in a single region is smaller. Second, and perhaps more importantly, the prediction here is for the peak infestation of the asymptotic wavefront. This definition makes sense mathematically to connect to species spread more generally (eg. Lewis et al. 2016; Kot 1992), but is not exactly equivalent to what we are observing. The initial condition for the forest in our simulation is set so that waves travel into forests where every tree is susceptible, and this allows the infestation wavefront to grow from its predicted equilibrium up to its peak. In reality, the forest will be more mixed age which would lead to a lower expected peak infestation (though managed stands or stands recovering from forest fire may have fairly similar ages (Safranyik and Wilson 2006)). In that case, though, it is harder to come up with a consistent definition for the size of an infestation. Importantly then, the size of the infestation should be interpreted as the fraction of susceptible trees that are infested, rather than the total fraction of pine density infested.

We note that our results suggest an effective management strategy may be to reduce the effective number of highly susceptible trees, in this case older and larger trees, and to additionally encourage a forest with a more mixed age structure. This strategy may be effective in reducing the overall number of beetles by reducing the absolute size of the wavefront, even if the fraction of susceptible trees infested remains high. Indeed, this strategy was adopted in the Alberta *Mountain Pine Beetle Management Strategy* (2007), where the stated goal was to reduce the number of highly susceptible stands by 25 percent over 20 years, and to change the age class structure over the landscape.

More broadly, our results here align with other studies assessing the risk of pine beetle spread under change. Using a phenomenological synthetic model, Cooke and Carroll (2017) consider the effects of warming, forest degradation, and host defense weakening and find similar results, while emphasizing future uncertainty of climate change. Other statistical models (eg. Sambaraju et al. 2012; Preisler et al. 2012; Xie 2024) similarly find that the range of MPB is likely to expand, but are often not able to directly address spread speed or size.

Our mechanistic model is able to provide insights into how predicted changes in forest health and beetle dynamics will translate into these metrics of spread. In particular, we find that MPB may spread more quickly and with greater severity under climate change. While we are not able to consider all possible factors that could lead to changes in model parameters, in particular beetle phenology, we focus on the parameter shifts most likely under climate change. This type of analysis is complementary to other studies, and allows us to further understand how MPB spread may change in the coming decades. While this specific model may not be directly applicable to other systems, similar mechanistic modelling may be able to address uncertainties in forest pest spread under climate change.

## Acknowledgments

We would like to thank all Lewis Research Group members, in particular Evan Johnson, Xiaoqi Xie, and Kévan Rastello, for their feedback on this project. We would also like to thank the members of the TRIA-FoR project for their support. MAL gratefully acknowledges the Gilbert and Betty Kennedy Chair in Mathematical Biology. Funding for this research has been provided through grants to the TRIA-FoR Project to MAL from Genome Canada (Project No. 18202) and the Government of Alberta through Genome Alberta (Grant No. L20TF), with contributions from the University of Alberta and fRI Research (Project No. U22004). This work was supported by Mitacs through the Mitacs Accelerate Program, in partnership with fRI Research. MB acknowledges the support of the Natural Sciences and Engineering Research Council of Canada (NSERC), [PDF – 568176 - 2022].

## Statements and Declarations

### Author contributions

Both authors contributed to study conception and design. The analysis was conducted by Micah Brush. The first draft of the manuscript was written by Micah Brush and both authors contributed to further edits.

